# Comparison of iSeq and Miseq in 16S rRNA sequencing-based human gut microbiome analysis

**DOI:** 10.1101/2025.08.22.671784

**Authors:** Yanling Wang, Vin Tangpricha, Andrew Gewirtz

## Abstract

Illumina’s MiSeq platform is a common approach in 16S-based microbiome analysis. Such usage is self-perpetuating in that many studies seek to employ widely used approaches to facilitate comparison of their results to existing literature. Yet, a range of factors, including cost and equipment availability can necessitate alternate approaches. For example, use of a Nano kit, lowers reagent costs by over 60% while others may only have access to entry-level sequencers such as Illumina’s iSeq. The extent to which these approaches would impact results and subsequently conclusions is unknown. Attempting to address this question from the literature is complicated in that various studies not only use distinct cohorts but also differ in the reagents/methodologies to used to isolate DNA and generate sequencing libraries. Hence, we sequenced a single 16S rRNA gene amplicon library derived from 60 fecal samples, collected during a dietary supplement intervention study via MiSeq, MiseqNano, and iSeq. We evaluated platform performance by several key measurements: alpha diversity, beta diversity, taxonomic composition, and differential taxonomic abundance analysis. We found that iSeq outperformed MiSeq-Nano in alpha diversity and differential abundance detection, while MiSeq-Nano provided better taxonomic resolution than iSeq. Most importantly, all platforms showed similar core biological patterns in alpha and beta diversity and overall taxonomic composition. Thus, MiSeq, MiseqNano, and iSeq are likely to yield the same biological conclusions although specific questions and logistical consideration may favor one of these approaches.

## Introduction

The advent of high-throughput sequencing technologies has led to the appreciation that gut microbiota composition is a major health determinant. While studies seeking deep mechanistic insights into how microbiota impacts health increasingly rely on shotgun metagenomic sequencing, 16s rRNA gene amplicon sequencing remains a workhorse of microbial analysis especially for studies seeking to track intra-individual changes over time in response to interventions (1).

The Illumina MiSeq platform is especially widely used for 16s sequencing, with the standard sequencing kit (2 × 250 bp, V2) generating approximately 20 million reads per run, resulting in a cost of around $1,500. For a standard 96-sample run, this provides an average sequencing depth of approximately 200,000 reads per sample. However, studies suggest that such deep sequencing may not always be necessary. For example, the Earth Microbiome Project rarefied sequences to just 5,000 reads per sample (2).

To provide more cost-effective alternatives, Illumina offers the Miseq-nano kit in Miseq system. It generates an average of 10,000 reads/sample for a standard 96 multiplexed run at a lower cost of approximately $400 per run. In addition, this Miseq-nano kit can offer same read length (2 × 250 bp) as the regular Miseq kit. However, the quality score of a Miseq-nano kit is usually lower than the full kit (see **Table 1**), due to only two tiles in the Miseq-nano flow cell and only top surface was imaged compared to 14 tiles and both top and bottom surface images in the full kit.

**Table 1.**
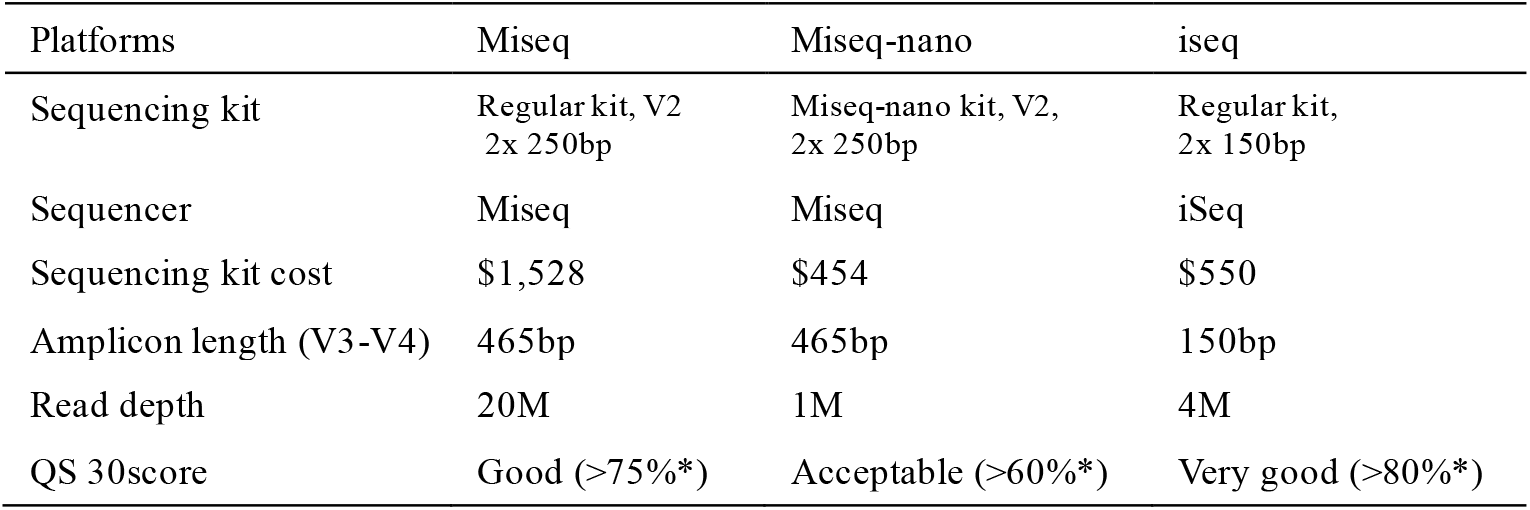
Characteristic comparison of three platforms.

Another alternative approach is the Illumina iSeq sequencer, released by Illumina in 2018 as a more affordable bench-top sequencer. The iSeq system itself costs significantly less (∼$20,000 vs. $100,000 for MiSeq). A regular iSeq kit generates approximately 40,000 reads/sample for a standard 96-well run, and the cost is only $500. A key limitation of the iSeq platform is its shorter read length, 2 × 150 bp. With joined paired-end reads, the longest amplicon iSeq can sequence is 280 bp (a minimum of 20-bp overlap is considered). However, the most targeted 16s rRNA gene regions are either V3–V4 (465bp) or V4 (291bp), both exceeding that capacity. As a result, 16s sequencing by iSeq sequencer often relies on single-end reads (150 bp), reducing taxonomic resolution compared to paired-end MiSeq sequencing (465bp for V3-V4 region).

Given the comparable cost of Miseq-nano and iSeq sequencing kit ($400–500 per run), researchers are faced with trade-offs: MiSeq-nano provides longer reads but fewer sequences and slightly lower quality scores, while iSeq offers higher quality scores and greater sequence depth but shorter reads. To evaluate the performance of these sequencing approaches relative to the standard Miseq kit, we conducted a comparative analysis using the same 16S rRNA gene library, prepared from 60 human fecal samples collected in a supplement intervention study (inulin and vitamin D) involving subjects with cystic fibrosis (CF) (3). The same 16s rRNA library was sequenced using three approaches: the standard MiSeq kit (hereafter referred to as “MiSeq”), MiSeq Nano kit (“Miseq-nano”), and iSeq (“iseq”). We compared the results in terms of alpha diversity, beta diversity, taxonomic composition, and differential abundance analysis to assess the strengths and limitations of each method for human gut microbiota research. Overall patterns of results were similar irrespective of sequencing methodology although some differences were noted.

## Results and discussion

### Alpha diversity

We first compared the gut microbiota diversity. Alpha diversity was evaluated by three metrics: the number of unique amplicon sequence variants (Observed Features), richness and evenness (Shannon), and Faith’s phylogenetic diversity (Faith PD). The differences in these metrics are summarized in **Table 2** (4).

**Table 2.** Alpha and beta diversity metrics.

**Tabel 2. Alpha and beta diversity metrics**
|  | Metrics | Qualitative or Quantitative | phylogenetic |
| --- | --- | --- | --- |
| Alpha diversity | Observed features | Qualitative | No |
|  | Shannon index | Quantitative | No |
|  | Faith phylogenetic diversity | Qualitative | Yes |
| Beta diversity | Jaccard | Qualitative | No |
|  | Bray Curtis | Quantitative | No |
|  | Unweighted unifracs | Qualitative | Yes |
|  | Weighted unifracs | Quantitative | Yes |

As shown in the rarefaction curve (**Figure 1A**), Miseq-nano detected the smallest number of Observed Features in these human stool samples, capturing only approximately a quarter of Observed Features detected in Miseq. iSeq performed better than Miseq-nano, capturing ∼50% of the Observed Features in Miseq. Faith PD follows a similar pattern to Observed Features. In contrast, the difference is minimal in the Shannon index across the three platforms, though Miseq-nano is still slightly lower than Miseq and iSeq. These results suggest that Observed Features and Faith PD are highly influenced by platforms, whereas Shannon remains relatively stable.

**Figure 1.**
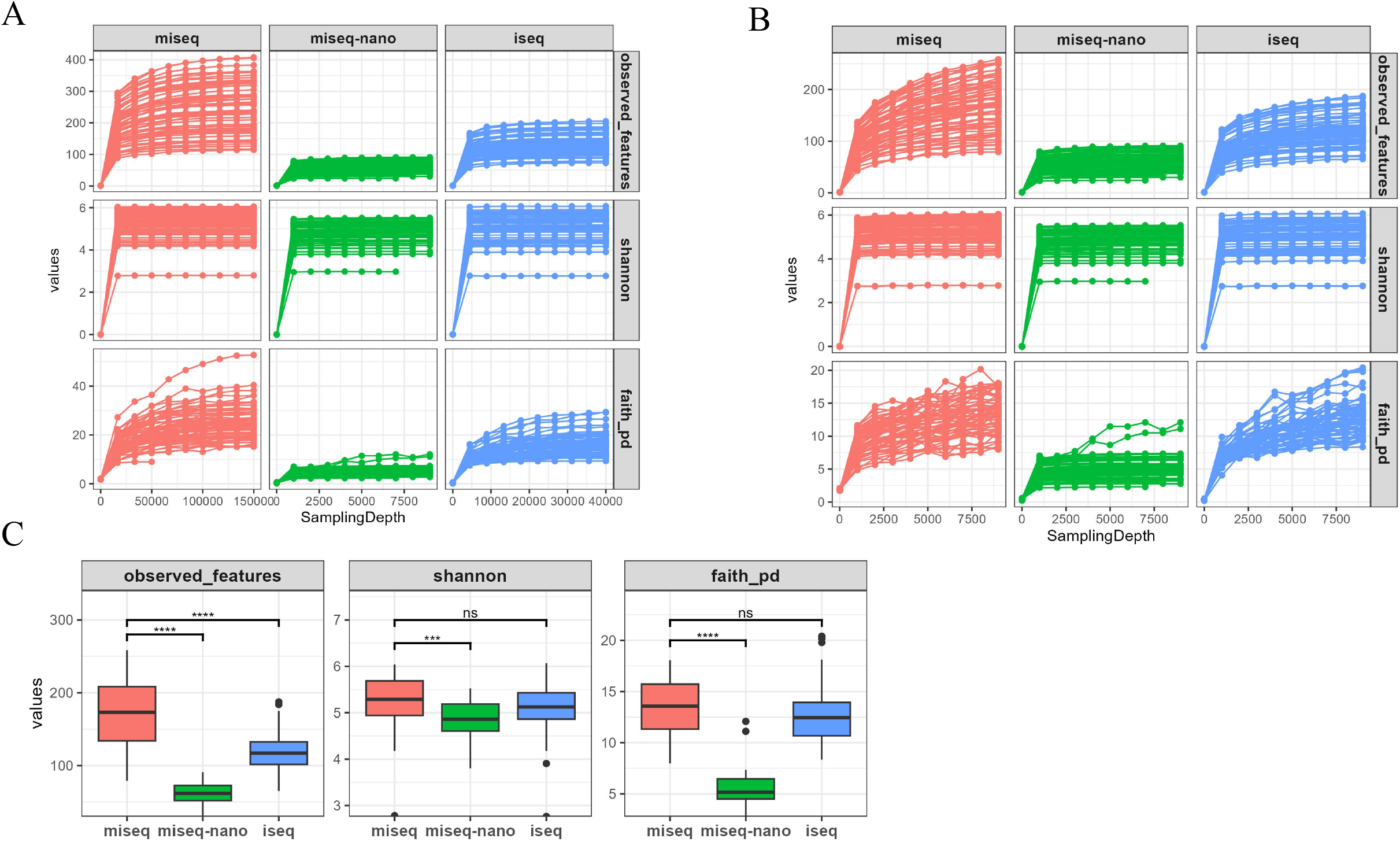
Overall distribution of alpha diversity across three platforms. (A-B) alpha rarefaction curve of each sample. (A) The maximum sampling depth was determined by half of the maximum reads for each platform: 150,000 reads for Miseq, 9,000 for Miseq-nano, 40,000 for iSeq. (B) Same sampling depth is selected for all platforms. (C) Box plot of alpha diversity at sampling depth of 9,000 reads. The Wilcoxon test was done to compare differences of alpha diversity, in reference to Miseq.

Unlike Observed Features and Faith PD, Shannon is a quantitative measure that places more weight on abundant features. For example, for a sample with 100 features, where 90 occur only once, is considered diverse in terms of Observed Features but not in terms of Shannon. Instead, a large Shannon index means the presence of many species that show well balanced abundances. The difference we observed between Shannon vs Observed Features, suggests that the additional features detected by MiSeq and iSeq are likely rare features.

Interestingly, even when rarefied at the same sampling depth (**Figures 1B and 1C**), Miseq-nano still exhibited significantly lower Observed Features and Faith PD than MiSeq, indicating that the differences between these two platforms may not be due to maximum sequencing depth alone, but rather intrinsic differences in flow cell design (e.g., there are only 2 tiles in Miseq-nano kit and only top surface is imaged whereas there are 14 tiles in standard Miseq kit, and both top and bottom surfaces are imaged).

As for iSeq, when rarefied at the same sampling depth, we found that the difference between iSeq and MiSeq became less pronounced (**Figures 1B and 1C**), and iSeq detected comparable levels of Faith and Shannon diversity as Miseq.

To probe the extent to which these platforms would detect intervention-induced changes, we analyzed the longitudinal alpha diversity changes by supplement treatments (Placebo, Inulin, VitaminD, VitaminD&Inulin) (**Figure 2**). We highlighted individual variation, as interpersonal differences are known to be substantial, often exceeding the effects of diet or lifestyle.

**Figure 2.**
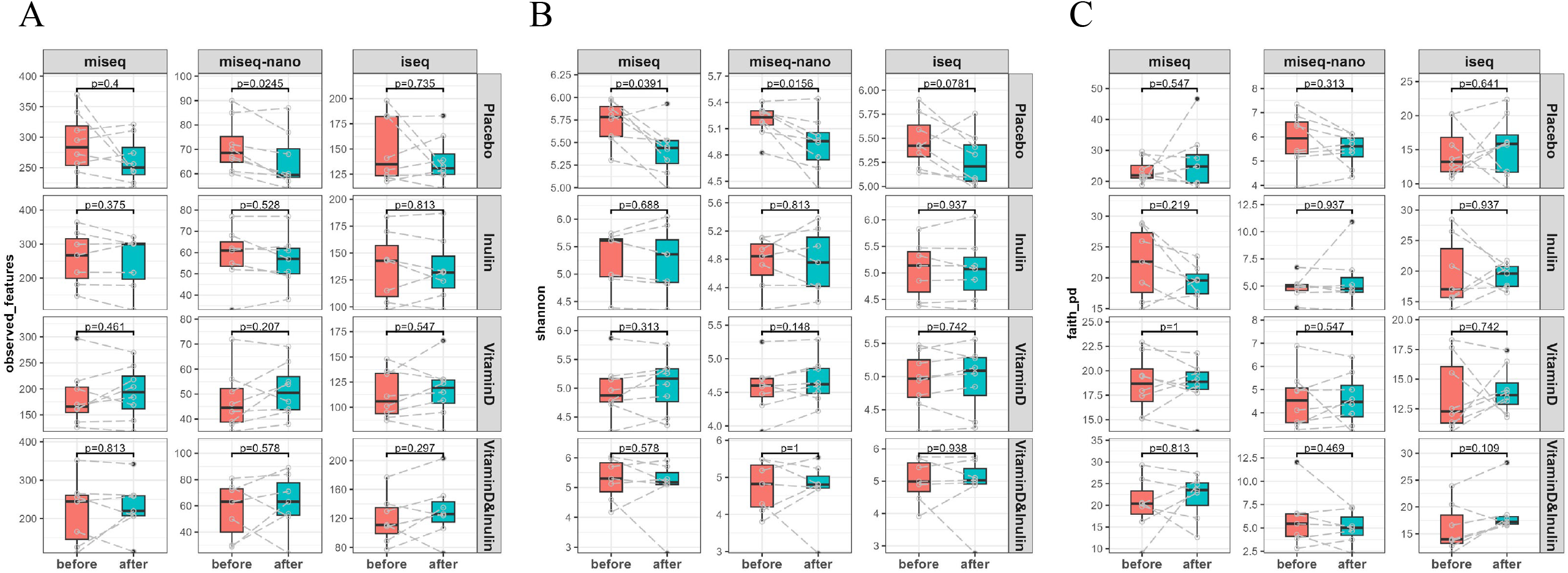
Longitudinal alpha diversity changes by supplements. Box plot of alpha diversity. Data from two timepoints (before and after) of each patient was connected by a line. Two-tailed paired Wilcoxon test was done to determine the difference between timepoints (p<0.05). (A) Observed Features. (B) Shannon. (C) Faith PD.

As shown in **Figure 2A**, despite notable differences in the absolute number of Observed Features across platforms, the treatment-associated relative changes in both MiSeq-nano and iSeq closely mirrors that observed with MiSeq. For instance, in the MiSeq dataset, most subjects exhibited a reduction in Observed Features following placebo treatment, and this trend that was successfully capitulated by both MiSeq-nano and iSeq, despite their lower absolute counts. A similar trend was observed in the other treatment groups. For the **Shannon index**, the absolute values were already comparable across the three platforms, and the treatment-associated relative changes followed similar patterns. Both MiSeq-nano and iSeq effectively captured the biological group differences observed in the MiSeq data (**Figure 2B**). In contrast, for **Faith’s Phylogenetic Diversity (Faith PD)**, the concordance among the three platforms was noticeably lower (**Figure 2C**).

Often, relative changes rather than absolute diversity values are more important for evaluating impacts of an intervention. In this case, when Observed Features or Shannon index were selected as the alpha diversity metrics, we found that both Miseq-nano and iSeq served as viable alternatives to MiSeq. However, if absolute alpha diversity numbers are desired, iSeq was a better option than Miseq-nano. Alternatively, Shannon, rather than other metrics, can be used as it is consistent across all three platforms, even with absolute numbers.

### Beta diversity

Determining whether the overall microbial community composition differs between treatment groups (beta diversity) is another important research question for 16s microbial sequencing. Given the substantial interpersonal variation noted earlier, we assessed beta diversity by measuring the community dissimilarity between timepoints within subjects. A larger beta diversity distance indicates a greater shift in gut microbial composition in these subjects following the supplement treatments. To provide a comprehensive analysis, we evaluated four beta diversity metrics: Jaccard, Bray-Curtis, Unweighted UniFrac, and Weighted UniFrac, with their differences summarized in **Table 2**. Like Faith PD in the alpha diversity, Unweighted and Weighted UniFrac incorporate phylogenetic information.

As shown in **Figure 3**, the patterns of longitudinal microbiota community changes across intervention groups were largely consistent across all three sequencing platforms. This was particularly evident for Bray-Curtis and Weighted UniFrac, the two quantitative beta diversity metrics that place more weight on abundant features. For example, in the MiSeq dataset, gut microbiota communities exhibited greater shifts following Placebo and Vitamin D & Inulin treatments compared to Inulin or Vitamin D alone, indicated by Weighted UniFrac distances, and this trend was reliably reproduced by both MiSeq-nano and iSeq (**Figure 3**, fourth row).

**Figure 3.**
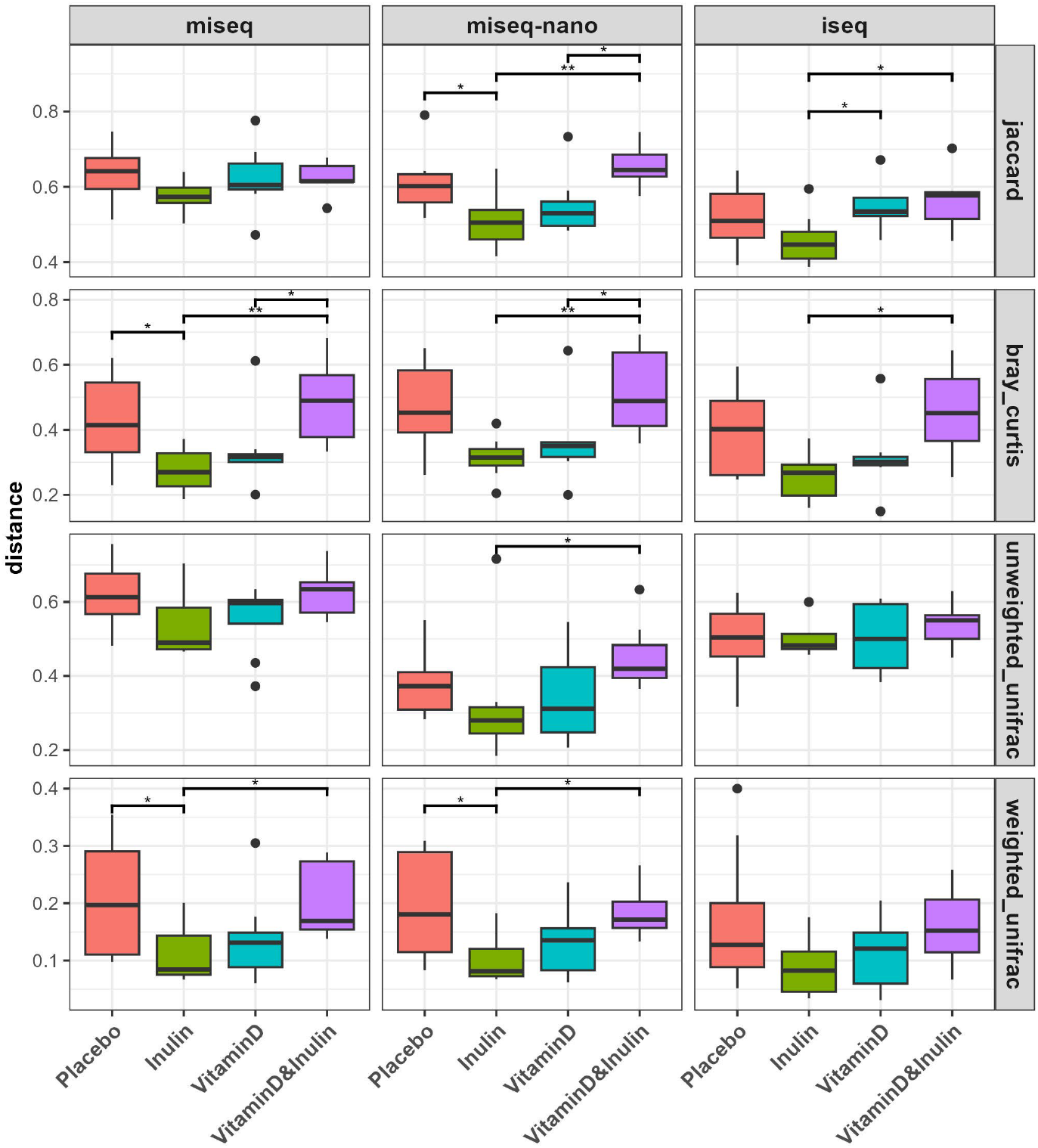
Microbiota community dissimilarity between timepoints. Box plot of four beta diversity distance metrics (Jaccard, Bray Curtis, Unweighted unifrac and Weighted unifrac). Two-tailed unpaired Wilcox test was done to determine the difference in supplement groups.

Nevertheless, MiSeq-nano more closely reproduced the statistical differences observed in the MiSeq data (**Figure 3**, fourth row).

In summary, both MiSeq-nano and iSeq effectively captured the beta diversity patterns identified by MiSeq, particularly for quantitative diversity metrics. However, MiSeq-nano demonstrated a slightly greater ability to replicate the statistical significance seen in the original MiSeq results.

### Taxonomic analysis

Another key objective of 16S sequencing is to characterize bacterial composition and identify taxa that differs among groups. At both phylum and genus levels, taxa distributions for Miseq-nano and iSeq closely resembled MiSeq, with Miseq-nano exhibiting slightly higher concordance with MiSeq (**Figures 4A, 4B**). We also noted that Miseq-nano, rather than iSeq, generates a more similar taxonomy profile to Miseq. As shown in **Figure 4C**, genera detected by Miseq are mostly overlapped with genera detected by Miseq-nano, while many genera detected by iSeq were only unique to iSeq.

**Figure 4.**
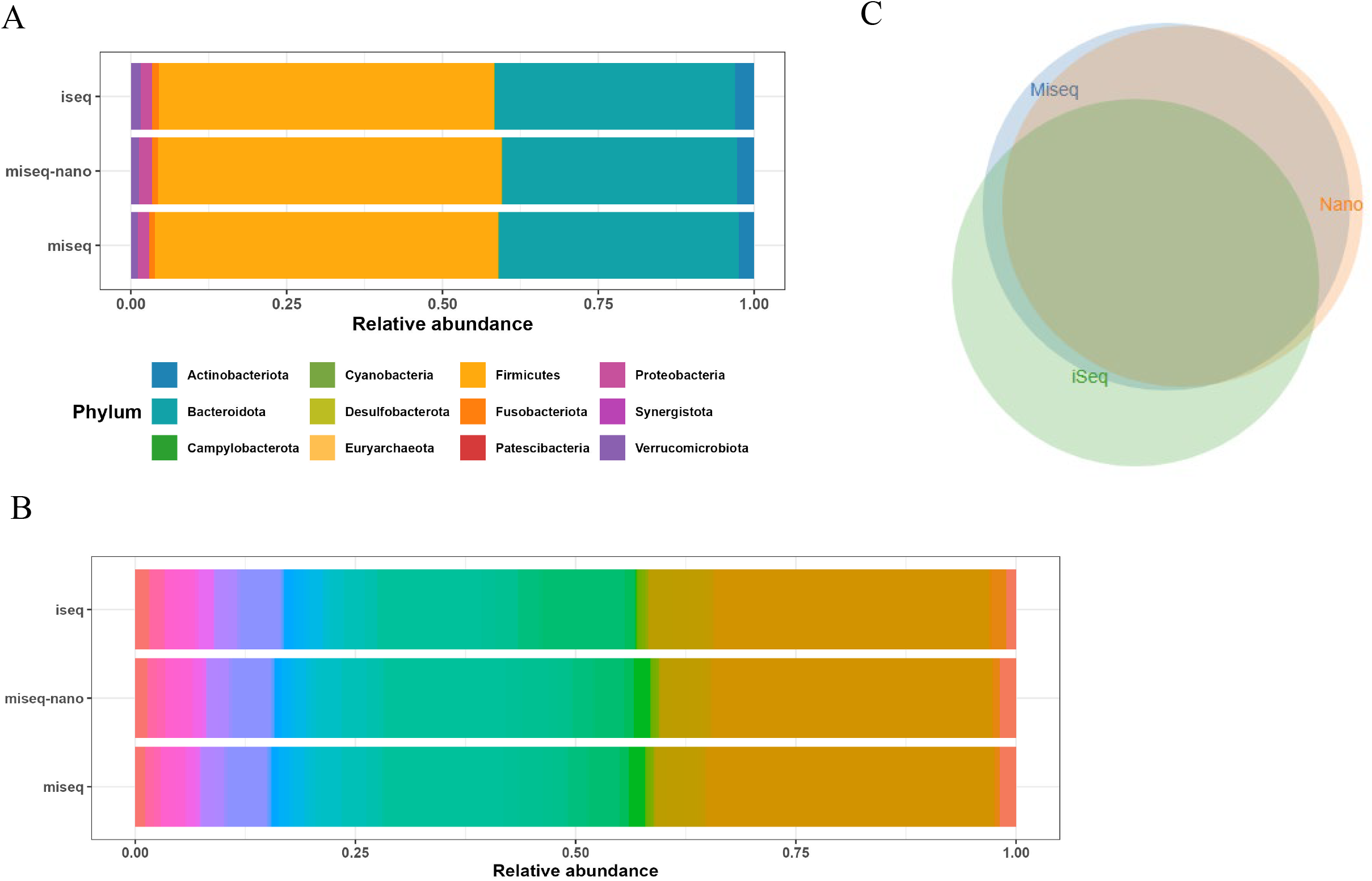
Relative taxonomy abundance by three platforms. (A) at phylum level. (B) at genus level. (C) Venn diagram of overlapping genera detected by the three platforms. Genera whose abundance are higher than 0.1% were selected. Blue circle represents genera detected by Miseq (N=58); orange by Miseq-nano (N=62) and green by iSeq (N=59).

For differential abundance analysis, we performed paired Wilcoxon tests to identify taxa with significantly different abundance between time points of each patient, for each intervention group. The results were visualized by a volcano plot (**Figure 5**), which summarizes both biological significance (fold change) and statistical significance (p-values). Taxa were considered significant if p < 0.05 and the fold change exceeded 2-fold.

**Figure 5.**
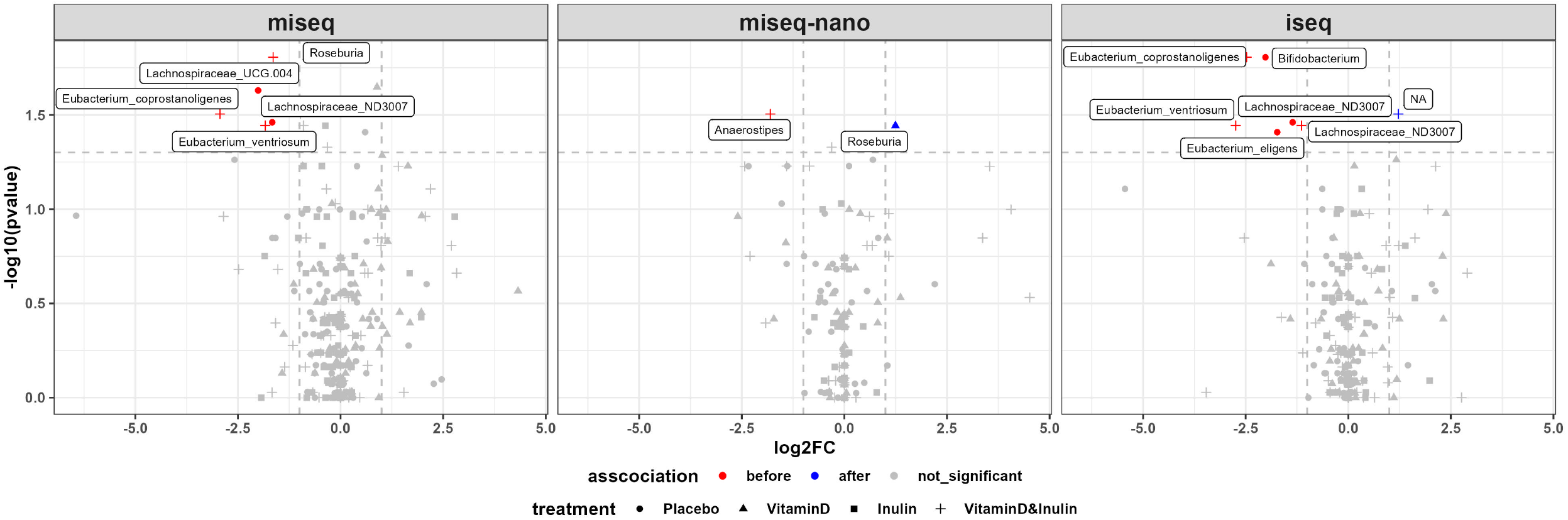
Taxonomy differential abundance analysis. Paired Wilcoxon tests were performed to detect changes in the abundance of each genus between time points (before and after the supplement intervention). Each dot represents a genus in each intervention group. Taxa were considered significant if p < 0.05 and the fold change exceeded 2-fold. Dots in red indicate genus which were significantly more abundant “before” the supplement intervention; dots in blue indicate genus which were more abundant “after” the supplement intervention; dots in grey indicate genus whose abundance were not significantly difference between time points. The shape of dots indicates intervention group. Names of significant genus are shown beside the dot.

As shown in **Figure 5**, MiSeq identified five significantly differential genus, Miseq-nano identified two, and Miseq-nano identified six. Specifically, Miseq identified genus *Lachnospiraceae_UCG*.*004* and *Lachnospiraceae _ND3007*, which were significantly more abundant before the Placebo intervention; genus *Roseburia, Eubacterium_coprostanoligenes*, and *Eubacterium_ventriosum* were significantly more abundant before the VitaminD& Inulin intervention. Among the five genera identified by Miseq, three were also identified by iSeq (*Lachnospiraceae_ND3007* before Placebo, and *Eubacterium_coprostanoligenes, Eubacterium_ventriosum* before VitaminD&Inulin), whereas none were identified by Miseq-nano. Additionally, Miseq-nano identified two false positives and iSeq two.

A closer examination of the five differential taxa identified by Miseq suggests that, although Miseq-nano did not identify any of them as statistically different, it still captured the trend of longitudinal changes after interventions (**Figure 6)**. For instance, genus *Eubacterium_coprostanoligenes* had a p-value of 0.0592 in Miseq-nano data, just above the cutoff, yet followed the decreasing trend closely matching that observed with Miseq (**Figure 6**, top row). Although Miseq-nano did not meet the statistical significance criteria, the corresponding biological pattern was still reflected in the data.

In summary, Miseq-nano generates a more similar taxonomy profile to Miseq than iSeq. For differential abundance analysis, iSeq demonstrates stronger statistical power and identifies more significantly different taxa. However, Miseq-nano should not be considered a complete failure— it still captures the biological trends, though without reaching significance. Researchers using Miseq-nano might consider adopting a more flexible p-value threshold to account for this limitation.

**Figure 6.**
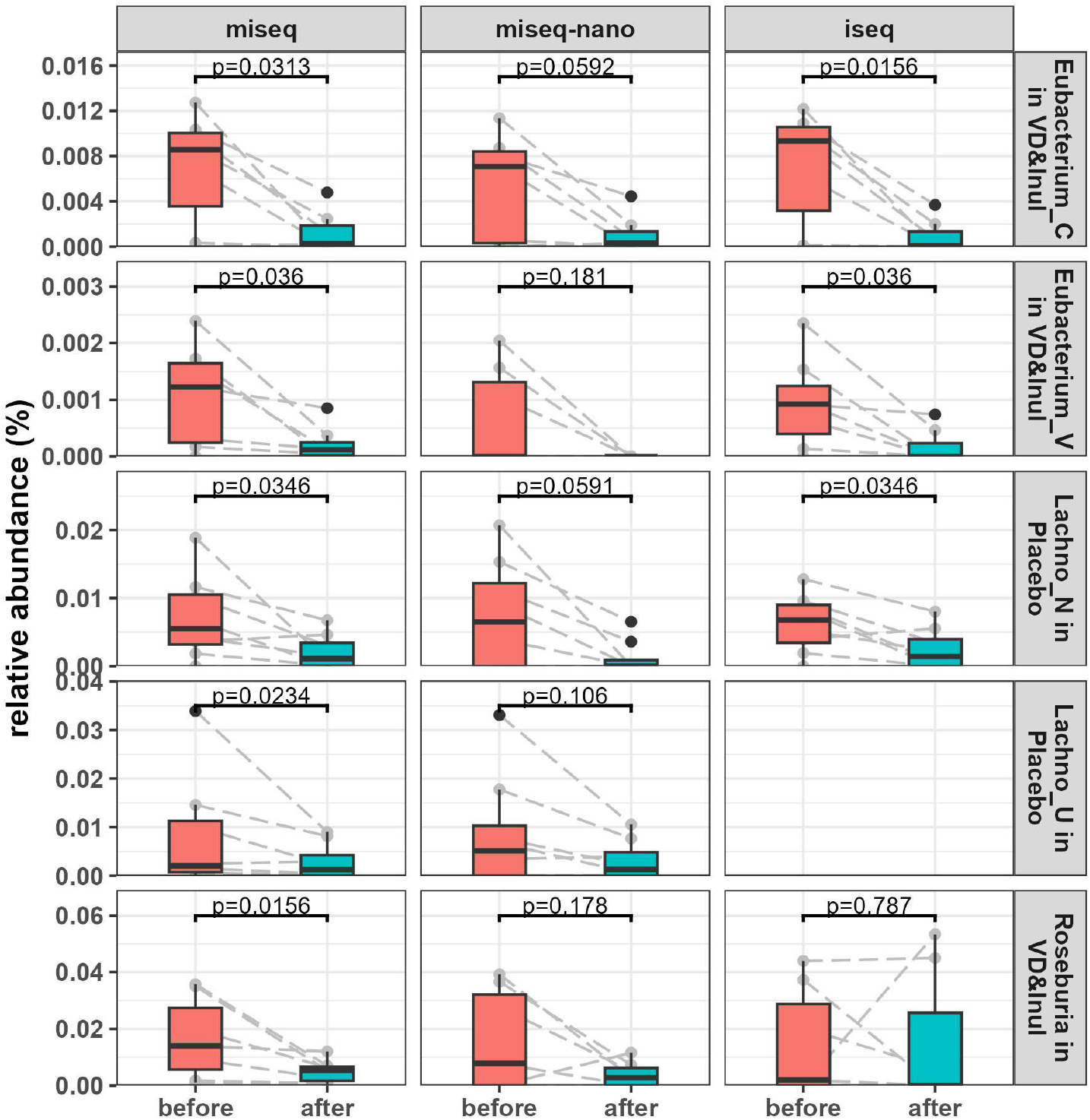
Relative abundance by three platforms of the five significant genera identified by Miseq.

## Conclusion

Our comparative analysis of MiSeq, MiSeq-nano, and iSeq platforms for 16S rRNA gene sequencing elucidated trade-offs between sequencing depth and read length but, nonetheless found that all three approaches are likely to yield generally similar overall conclusions. In reference to the standard Miseq method, both Miseq-nano and iSeq offer cost-effective alternatives suitable for many specific research goals. iSeq, which has a much lower instrument cost but a higher reagent cost, outperformed Miseq-nano in terms of alpha diversity, and differential abundance analysis, likely due to its higher sequencing depth and quality scores. However, for targeted taxa research, Miseq-nano outperforms iSeq in taxa identification, likely due to its longer reads. Both Miseq-nano and iSeq preserved key biological patterns across samples observed in the standard MiSeq approach, especially in beta diversity and overall taxonomic composition. Ultimately, the optimal choice of sequencing method should be guided by study objectives, sample nature and target amplicon length. A limitation of this study is that the extent to which our conclusions apply to other samples remain unknown. Indeed, optimal sequencing strategies still depend on the nature of the sample and research objectives.

## Methods

### Samples

The 16s rRNA library was obtained from the study (3). The study was a 2 × 2 factorial, double-blinded, placebo-controlled, randomized, pilot clinical trial in which 30 CF subjects received oral cholecalciferol (vitamin D3) (50,000 IU/week) and/or inulin (12 g/day) for 12 weeks. Thus, there were 4 study groups: placebo (n = 8), vitamin D (n = 8), inulin (n = 7), vitamin D & inulin (n = 7). 60 stool samples were collected at baseline (just before) and after the intervention. Fecal DNA was extracted from the samples, and V3-V4 region of 16S rRNA genes were amplified using the following primers: 341F 5′ TCGTCGGCAGCGTCAGATGTGTATAAGAGACAGCCTACGGGNGGCWGCAG-3′; 805R 5′ GTCTCGTGGGCTCGGAGATGTGTATAAGAGACAGGACTACHVGGGTATCTAATCC-3′.

After index PCR and purification, an equal molar of each sample was then combined as the library. The same library was then sequenced on Miseq system with the standard kit Miseq V2, 2 × 250 bp (hereafter referred to as “MiSeq”), Miseq system with Miseq-nano kit V2, 2 × 250 bp (“Miseq-nano”), iSeq system with iSeq kit, 2 × 150 bp (“iSeq”).

## Data analysis

Demultiplexed fastq files were generated on instrument and imported in Qiime2 (version Qiime2-amplicon-2023.09) for analysis (5). For Miseq and Miseq-nano analysis, paired end reads were joined. For iSeq analysis, only read 1 was analyzed. Low quality scored reads were filtered by DADA2 plugin (6). After DADA2, a range of 55,174–293,372 reads (median 182,419) per sample was obtained for Miseq. A range of 4,737-17,520 reads (median 11,209) per sample was obtained for Miseq-nano. A range of 33,532-75,032 reads (median 49,828) per sample was obtained for iSeq.

For alpha and beta diversity analysis, reads were rarefied to 55,174 reads per sample for Miseq, 4,737 for Miseq-nano and 33,532 for iSeq. Alpha and beta diversity metrics were computed by q2-diversity plugin. A rooted phylogenetic tree was generated by align-to-tree-mafft-fasttree pipeline from the q2-phylogeny plugin (7, 8). The rooted tree was used to compute phylogenetic diversity metrics, including Faith’s Phylogenetic Diversity (Faith PD), weighted and unweighted UniFrac.

Taxonomy was assigned to ASVs using the q2-feature-classifier (9) classify-sklearn naïve Bayes taxonomy classifier against bespoke Silva 138.1 classifier, animal distal gut habit (10), house trained for V3-V4 region. For differential abundance analysis, features were collapsed at genus level. After normalization (converting the counts into relative abundances), a pseudo count (nine-tenth of the lowest non-zero value in the relative abundance table) was added to all the data. Two-tailed paired Wilcoxon tests were performed to detect changes in the abundance of each taxa between time points (before and after the supplement intervention). As the primary focus is to compare the three platforms, p-values shown in this paper are all unadjusted. Media fold change was used to represent the fold change of each taxa before vs after. P-value less than 0.05 and fold change larger than 2-fold were considered as significant.

All statistical tests and plots were done in R version 4.2.3.

## Acknowledgements

This study was supported by National Institutes of Health grants DK083890 and DK099071 to A.T.G; P30DK125013 and 3UL1TR002378-05S2 to V.T.; Cystic Fibrosis Foundation Awards TANGPR19A0 and CC002-AD to V.T.

---

**Supplement Figure 1.**
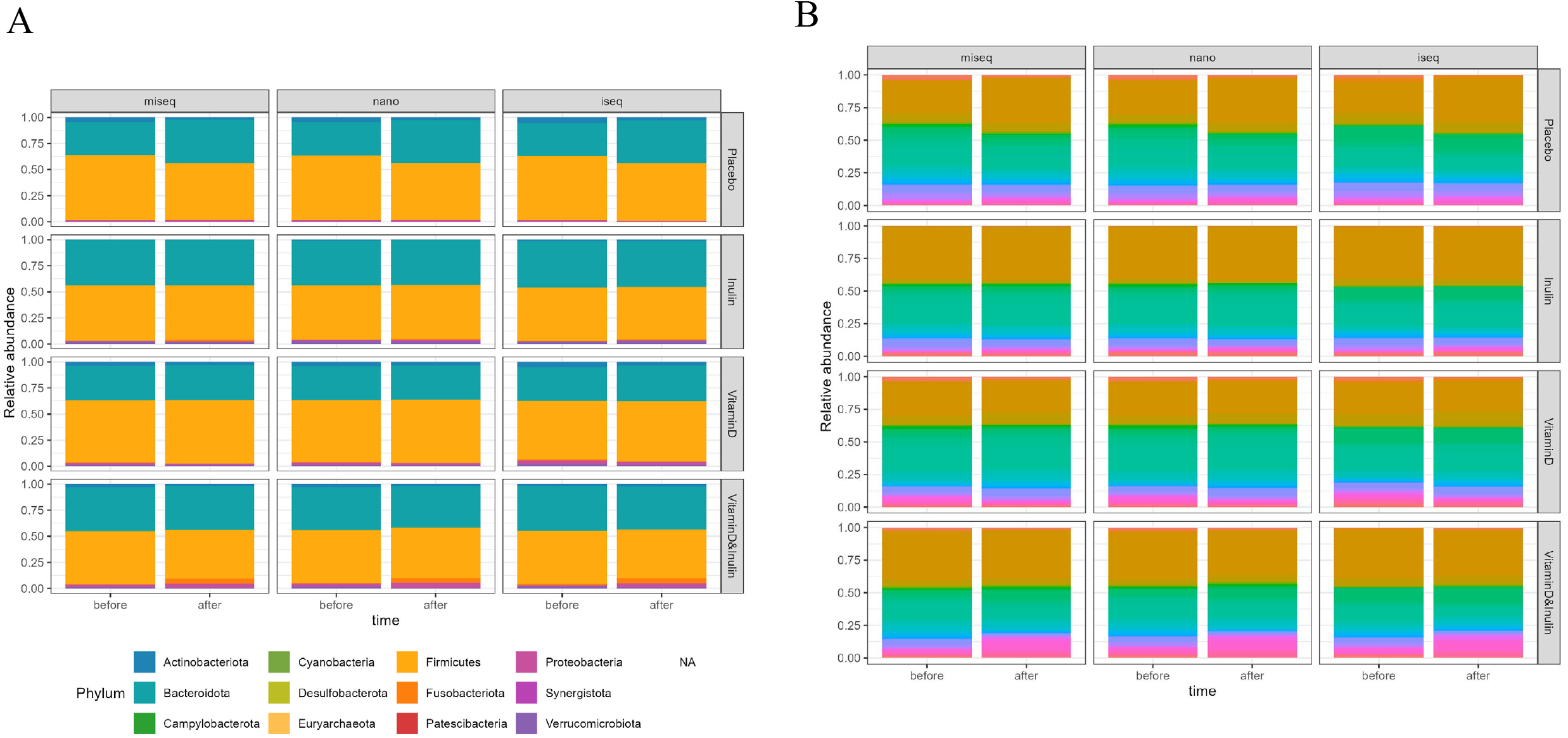

## Notes

### Competing Interest Statement

The authors have declared no competing interest.

